# PLK4 is a microtubule-associated protein that self assembles promoting *de novo* MTOC formation

**DOI:** 10.1101/312272

**Authors:** Susana Montenegro Gouveia, Sihem Zitouni, Dong Kong, Paulo Duarte, Beatriz Ferreira Gomes, Ana Laura Sousa, Erin M. Tranfield, Antony Hyman, Jadranka Loncarek, Monica Bettencourt-Dias

## Abstract

PLK4 binds to microtubules and self assembles into supramolecular assemblies that recruit tubulin and trigger *de novo* MTOC formation in *Xenopus laevis* extracts.

**Abstract:** The centrosome is an important microtubule-organizing center (MTOCs) in animal cells and it consists of two barrel-shaped structures (centrioles), surrounded by the pericentriolar material (PCM), which nucleates microtubules. PCM components form condensates, supramolecular assemblies that concentrate microtubule nucleators. Centrosomes can form close to an existing structure (canonical duplication) or *de novo*. How centrosomes form de novo is not known. PLK4 is a master driver of centrosome biogenesis, which is critical to recruit several centriole components. Here, we investigate the beginning of centrosome biogenesis, taking advantage of *Xenopus egg* extracts, where we and others have shown that PLK4 can induce *de novo* MTOC formation (Eckerdt et al., 2011; Zitouni et al., 2016). Surprisingly, we observe that *in vitro,* PLK4 can self-assemble into supramolecular assemblies that recruit α/β-tubulin. In *Xenopus* extracts, PLK4 supramolecular assemblies additionally recruit the PLK4 substrate STIL and the microtubule nucleator, γ-tubulin, and form acentriolar MTOCs *de novo*. The assembly of these robust microtubule asters is independent of dynein, similarly to centrosomes. We suggest a new mechanism of action for PLK4, where it forms a self-organizing catalytic scaffold that recruits centriole components, PCM factors and α/β-tubulin, leading to MTOC formation.

## Introduction

Centrosomes are important microtubule organizing centres (MTOCs) in animal cells, being involved in a variety of cellular and developmental processes, including cell motility, division and polarity (Sanchez and Feldman, 2017). Centrosomes are composed of a core structure, a pair of centrioles, surrounded by a protein-rich pericentriolar material (PCM), which nucleates and anchors a microtubule (MT) array within the cell (Paz and Luders, 2017). PCM proteins can also associate with other cellular structures to assemble non-centrosomal MTOCs (Sanchez and Feldman, 2017), the assembly of which is less characterised (Sanchez and Feldman, 2017).

Critical for centrosome assembly is PLK4, a serine-threonine kinase, member of the polo-like kinase family, which triggers procentriole formation close to a centriole that already exists, or induces centriole *de novo* formation when centrioles are absent (Bettencourt-Dias et al., 2005; Habedanck et al., 2005; Rodrigues-Martins et al., 2007). Recently, it was demonstrated that PLK4 promotes MT nucleation in the acentriolar mouse embryo, being essential for spindle assembly, suggesting it can also contribute to acentriolar MTOC formation (Bury et al., 2017; Coelho et al., 2013).

How PLK4 protein drives de novo MTOC formation is not understood. To study the role of PLK4 in acentriolar systems, we used both *in vitro* systems and acentriolar *Xenopus* extracts, where it had been previously observed that PLK4 is sufficient to generate *de novo* MTOCs (Eckerdt et al., 2011; Zitouni et al., 2016). We show that *in vitro* PLK4 self-assembles into supramolecular assemblies that recruit tubulin. In *Xenopus* extracts, PLK4 supramolecular assemblies recruit STIL, γ-tubulin and tubulin, forming acentrosomal MTOCs *de novo*. Thus, PLK4 plays an important role in forming both centriole-containing and acentriolar MTOCs.

## Results and Discussion

### PLK4 self-assembles into supramolecular assemblies that concentrate soluble tubulin *in vitro*

We wished to investigate how PLK4 drives de novo MTOC formation. Recently, Woodruff and colleagues used a minimal set of *C. elegans* proteins to reconstitute a functional MTOC *in vitro*. They observed that the PCM protein, SPD-5, self-assembles into spherical scaffolds named condensates, which together with homologs of XMAP215 and TPX2 allowed the formation of acentrosomal MTOCs (Woodruff et al., 2017). SPD-5 condensates are formed in macro-molecular crowding environments containing polyethylene glycol (PEG) and once formed can concentrate other proteins (Woodruff et al., 2017).

We wished to also explore a minimal system to study PLK4 function. We expressed GFP tagged *Xenopus* PLK4 in the baculovirus system. We were surprised to observe that purified GFP-PLK4, but not GFP alone, self-assembles into sphere-like structures similar to the spherical SPD-5 condensates, in this case even in the absence of PEG (Fig. 1A, 1B) (Woodruff et al., 2017). It is possible that PLK4 supramolecular assemblies form through multimerization, as PLK4 has the ability to dimerise in different regions of the protein (Jana et al., 2014).

**Fig. 1.**
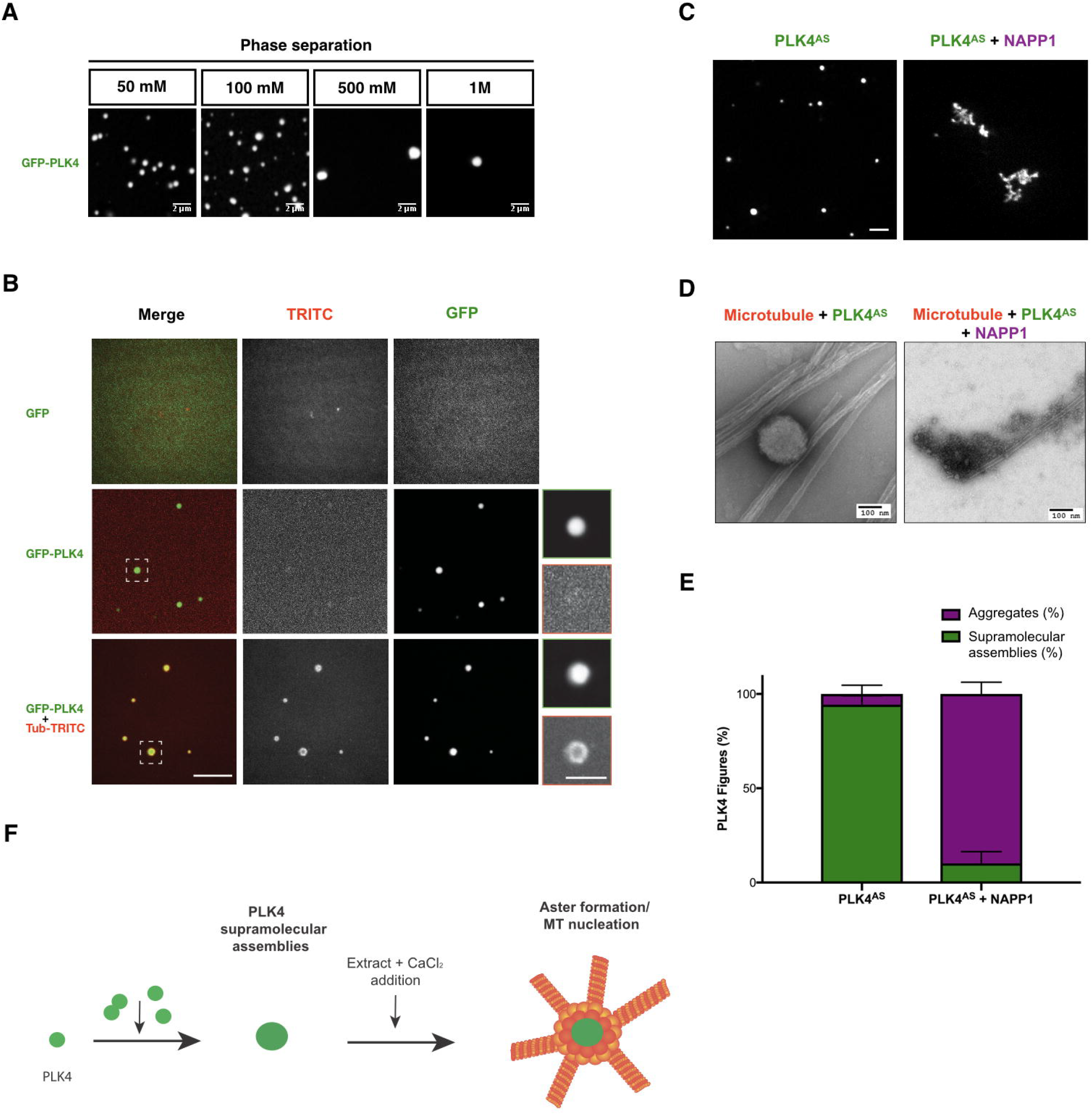
PLK4 self-assembles into supramolecular assemblies that concentrate tubulin *in vitro* and is dependent on its kinase activity. (A) Representative images of GFP-PLK4 supramolecular assemblies formed at different concentrations of NaCl. (B) Representative confocal images of GFP-PLK4 condensate formation in the absence or presence of rhodamine-labelled tubulin (500 nM). GFP was used as a control. Scale bars: 5 μm; Insets: 2 μm. (C) Confocal images representing GFP-PLK4^AS^ in the absence or presence of NAPP1. Scale bars: 5 μm. Note that in presence of NAPP1, GFP-PLK4 forms disorganized structures. (D) Electron microscopy (EM) images of GFP-PLK4^AS^ in presence or absence of NAPP1. Scale bars: 100 nm. (E) Quantification of percentage of supramolecular assemblies versus aggregates obtained from EM data. Three independent experiments were counted. (F) Scheme illustrating the results.

We then asked whether PLK4 on its own would form an MTOC, and added α/β-tubulin to the assay. Using confocal microscopy, we observed that PLK4 supramolecular assemblies are able to selectively recruit α/β-tubulin (Fig. 1B). Although we were unable to observe MT nucleation *in vitro*, we were intrigued by the fact that α/β-tubulin coats PLK4 supramolecular assemblies without the need for MT nucleators (Fig. 1B).

PLK4’s ability to form centrioles in cells and MTOCs in *Xenopus* extract requires its kinase activity (Rodrigues-Martins et al., 2007; Zitouni et al., 2016). We asked whether supramolecular assembly requires PLK4 kinase activity, using recombinant GFP-PLK4^AS^ (L89A/H188Y). PLK4^AS^ can specifically fit bulky ATP analogues, making it sensitive to ATP-analogue inhibitors such as NAPP1, while having a comparable kinase activity to PLK4^WT^ (Bishop et al., 2000; Zitouni et al., 2016). We show that recombinant GFP-PLK4^AS^ has the ability to self-assemble into supramolecular assemblies similar to GFP-PLK4^WT^ (Fig. 1C and 1D). However, in the presence of the inhibitor NAPP1, the formation of supramolecular assemblies is severely impaired, as observed by confocal microscopy (Fig. 1C) and electron microscopy (Fig. 1D and 1E). Instead of robust spherical structures, PLK4 aggregates show an amorphous network with no regular shape or higher-order structure (Fig. 1D). We observed the same effect when PLK4 was treated with lambda phosphatase, which renders it inactive, suggesting that the catalytic activity of PLK4 is required to promote the formation of supramolecular assemblies *in vitro* (Fig. S1B) (Lopes et al., 2015).

### PLK4 binds microtubules *in vitro*

Given that PLK4 supramolecular assemblies can recruit tubulin, we asked whether PLK4 has affinity for MTs and could promote MT stabilization. We observed by confocal microscopy that GFP-PLK4 supramolecular assemblies associate with stable MT seeds *in vitro* when they are incubated together (Fig. 2A). The great majority of PLK4 supramolecular assemblies are associated to MTs (~95.4%) (Fig. 2B). We then asked if PLK4 binds directly to MTs. We performed MT pelleting assays, where PLK4 and MTs are ultracentrifuged together. Polymerized MT will pellet (P) together with bound protein whereas the unbound fraction will remain suspended in solution (S). We observed that purified PLK4 is able to co-pellet with the MT fraction *in vitro* (Fig. 2C). To calculate the binding dissociation constant (Kd), we performed pelleting assays with a constant concentration of PLK4 (0.7 μM) and increasing concentrations of MTs (0 to 4 μM). Reciprocally, we performed the same assay using a constant amount of MTs (10 μM) and increasing amounts of PLK4 (0 to 4 μM) until the saturation point, (Fig. 2C). We further plotted PLK4 bound to MTs versus MT concentration, from three independent experiments. The calculated Kd is the concentration of MTs that is required to sediment half of PLK4 (Fig 2D). These data strongly indicate that PLK4 is a MT-associated protein (MAP) that binds MTs directly with high affinity (Kd= 0.62 μM ± 0.071). In addition, PLK4 kinase activity does not seem to be required for MT binding since the inhibited recombinant GFP-PLK4^AS^ still binds MTs (Fig 1D). Finally, we observed that PLK4 led to an increase in the formation of MT bundles in a concentration-dependent manner (Fig. 2E and 2F). As MT bundles are known to stabilize MTs dynamic, perhaps PLK4 promotes MT stabilization (Brandt and Lee, 1994; Umeyama et al., 1993).

**Fig. 2.**
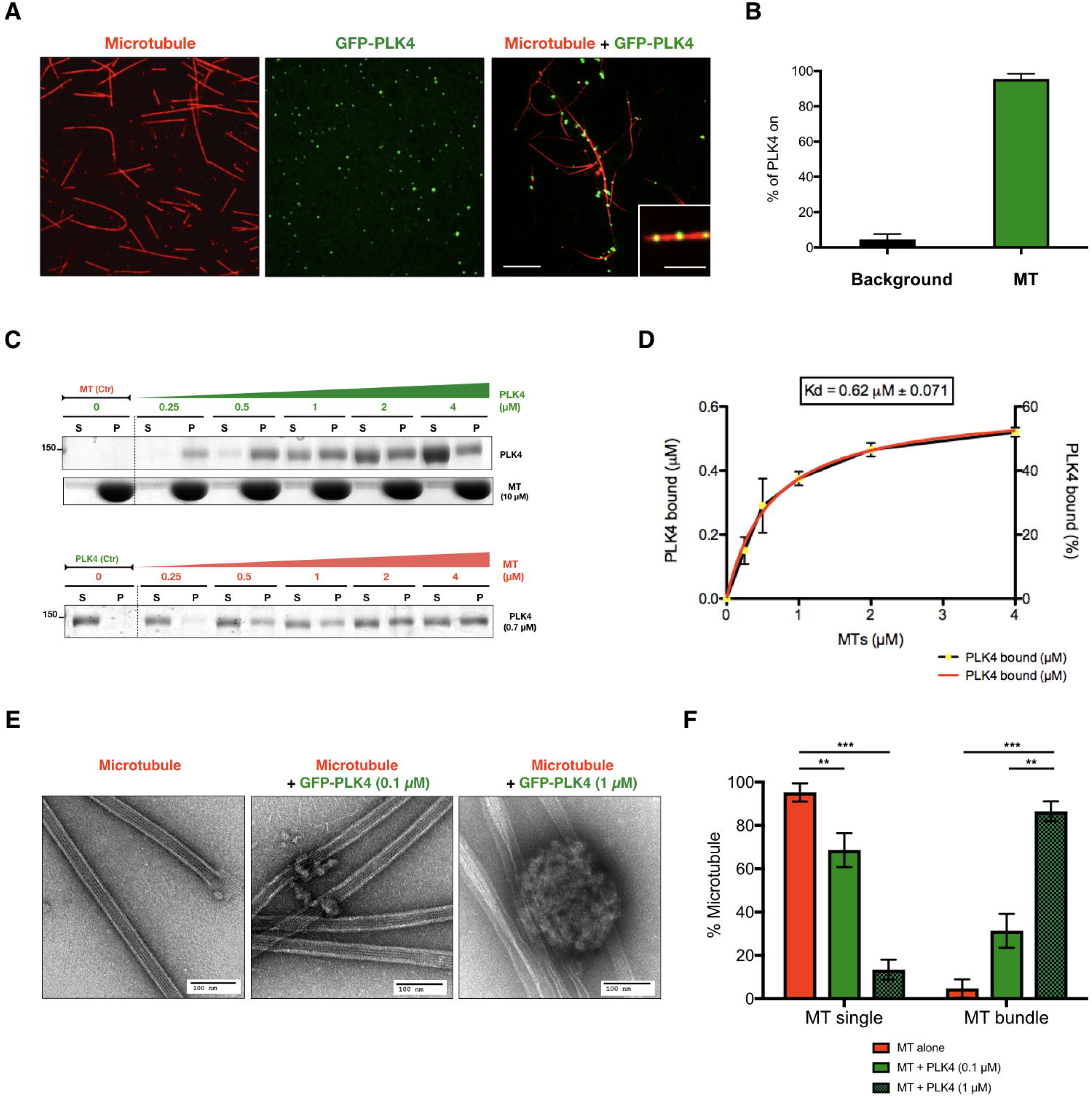
PLK4 is a microtubule-associated protein that promotes microtubule bundling *in vitro*. (A) Confocal images of Taxol-stabilized MTs alone (rhodamine-labeled tubulin, red), recombinant purified GFP-PLK4 alone (green) and the mixture of both. Scale bar: 5 μM; Inset: 2 μm. (B) Quantification of PLK4 supramolecular assemblies associated to MTs compared to free PLK4 supramolecular assemblies in the background. (N=3, n=100 spot/conditions). (C) MT-pelleting assays. The two assays are showing a constant concentration of PLK4 (0.7 μM) mixed and incubated with different concentrations of MTs (0 to 4 μM) or an increasing amounts of GFP-PLK4 (0 to 4 μM) in presence of constant concentration of MTs (10 μM). The western blot is showing supernatant (S) and pellet (P) for each condition. (D) Quantitative analysis of the binding properties between PLK4 and MTs. Note that the dissociation constant (Kd) for PLK4, determined by best fit to the data (red curve), is 0.62 ± 0.071 μM. Data were collected from three independent experiments. (E) EM images showing MTs alone or MTs incubated with two different concentrations of PLK4 (0.1 μM and 1 μM). Scale bars: 100 nm. (F) Percentage of single or bundled MTs quantified from the EM data in the presence of PLK4 at 0.1 μM or 1 μM; MTs alone are used as a control. Results were scored using 30 images per condition obtained from 3 independent experiments each; (***P<0.001; **P<0.05).

### PLK4 supramolecular assemblies form *de novo* MTOCs in *Xenopus* extracts that mimic centrosomes *in vivo*

We asked whether PLK4 supramolecular assemblies could promote MT nucleation when exposed to the right environment. It was previously shown that PLK4 induces *de novo* MTOC formation after exit from M-phase in *Xenopus egg* extracts (Eckerdt et al., 2011; Zitouni et al., 2016). We observed that PLK4 supramolecular assemblies, after being formed *in vitro*, are able to nucleate MTs if incubated with *Xenopus egg* extracts, suggesting that PLK4 supramolecular assemblies act as a scaffold that forms an active MTOC (Fig. 3A).

**Fig. 3.**
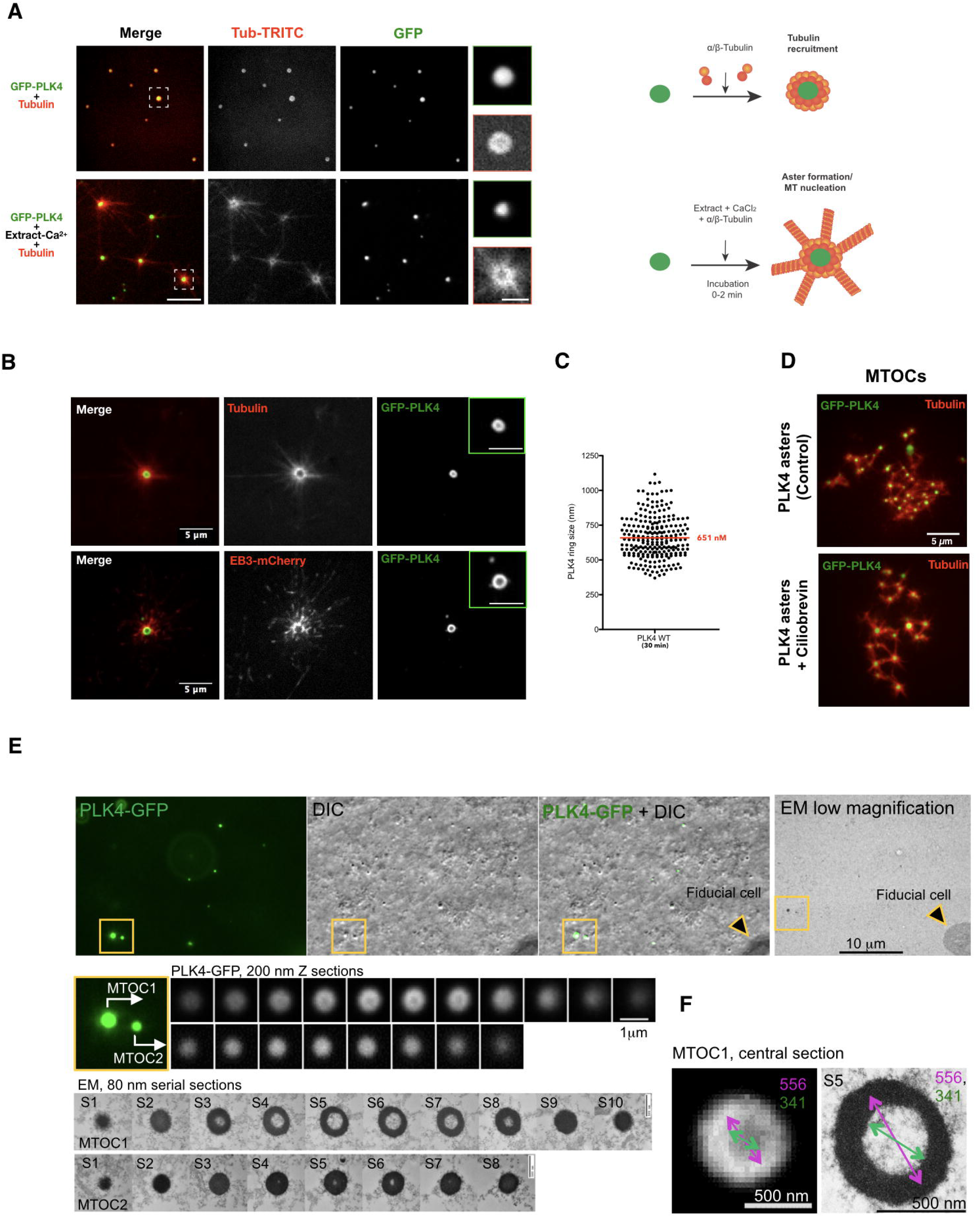
PLK4 supramolecular assemblies form *de novo* MTOCs in *Xenopus* extracts that are independent of motor proteins and mimic centrosomes *in vivo*. (A) GFP-PLK4 was mixed concomitantly with rhodamine-labeled tubulin or supramolecular assemblies were formed (step I) and then extract released to interphase with calcium containing rhodamine-labeled tubulin were added to these supramolecular assemblies (step II). Note that nucleation was observed instantly after the addition of the mixture (0-2 min). Scale bars: 5 μm, inset= 2 μm. (B) Confocal images showing MTOC formation in *Xenopus* Mll-calcium-released extracts in presence of recombinant GFP-PLK4 (green). MTs are visualized using rhodamine-labelled tubulin (red) (upper panel) and EB3-mCherry (lower panel). MT plus ends visualized by EB3-mCherry point out to the edge of the aster. The insets show PLK4 as a ring-like structure (see Movie 1). (C) Quantification of the size (nm) of GFP-PLK4 ring-like structure after 30 min of incubation. GFP-PLK4 rings were measured from 3 independent experiments. (D) PLK4 asters are independent of dynein. Representative confocal images of PLK4 asters are shown in the control and in the presence of ciliobrevin (dynein inhibitor). (E) (F) Correlative light/electron microscopy analysis of PLK4’s MTOCs. Plk4-GFP signals were first visualized by fluorescence and DIC, and then by electron microscopy. A series of 200 nm sections (confocal) and 80 nm EM sections are presented for two MTOCs (yellow box, MTOC1 and MTOC2) (G) Measurememts of the central sections of MTOC1 (section S5 in F). Values (nm) are presented in the table. Scale bars: 500 nm and 1000 nm.

Next we investigated whether GFP-PLK4 forms supramolecular assemblies in extracts. Supramolecular assemblies were formed in the extract and were variable in size (Fig. 3C), with sizes similar to the size of the centrosome (300 to 1000 nm, with an average size of ~650 nm). We used rhodamine-tubulin and EB3-mCherry to visualize the nucleation driven by GFP-PLK4 in extracts (Fig. 3B, Movie 1). These MTOCs contain GFP-PLK4 at their core, showing a ring-like-structure, surrounded by tubulin, as observed by confocal microscopy. This is similar to what we observed *in vitro*, suggesting we are looking at the same entity both *in vitro* and in the extract.

Microtubule asters can be formed in two different ways: motor based self-assembly of MT minus-end bound material (acentrosomal MTOCs) (Compton, 1998; Mitchison, 1992; Sanchez and Feldman, 2017), where motor proteins, such as dynein, play a crucial role in MTOC formation (Gaglio et al., 1997; Gaglio et al., 1996); or alternatively in a motor-independent manner, relying on nucleation and anchoring of MTs to a pre-existing structure such as the centrosome. We thus investigated whether PLK4 MTOCs depend or not on dynein. Centrioles and DMSO asters were used as controls. As expected, while DMSO asters are destroyed in the presence of the dynein inhibitors, vanadate and ciliobrevin, centrioles remained capable of nucleating MTs (Fig. S2). In the case of PLK4 driven MTOCs, we observed they could form in a dynein-independent manner (Fig. 3D and S2), showing their independence from motors. This suggests PLK4-driven MTOCs form in a similar manner to centrosomes.

We then investigated whether there were centrioles at the center of the aster, investigating the ultrastructure of PLK4-driven MTOCs, using correlative light-electron microscopy (CLEM) (Fig. 3E). Unexpectedly, we observed no centrioles. Instead, we observed sphere like structures; these structures are correlated with the ring structure we observed by confocal microscopy and are the centre of PLK4 MTOCs. Most of the structures were hollow (Fig. 3F), with some exceptions; their size was on average ~700 nm (Fig. 3E, 3F and 3G). We conclude that PLK4 supramolecular assemblies localize at the centre of the PLK4-induced MTOCs and have the ability to nucleate MTs, similar to bona-fide centrosomes.

### PLK4 supramolecular assemblies recruit STIL and γ-tubulin in *Xenopus* extracts, leading to centrosomal MT nucleation

To further characterize PLK4 supramolecular assemblies, and understand their ability to form an MTOC, we asked whether these supramolecular assemblies also recruit other components (in addition to α/β-tubulin), in particular PLK4 substrates and MT nucleators. First, we used 3D-SIM to characterize at super-resolution level PLK4-driven MTOCs in *Xenopus* egg extracts. We could observe the GFP-PLK4 structure similar to a ring in its centre in 2D (Fig. 4A) and to a sphere/condensate after 3D reconstruction (Fig. 4B and Movie 2 and 3).

**Fig. 4.**
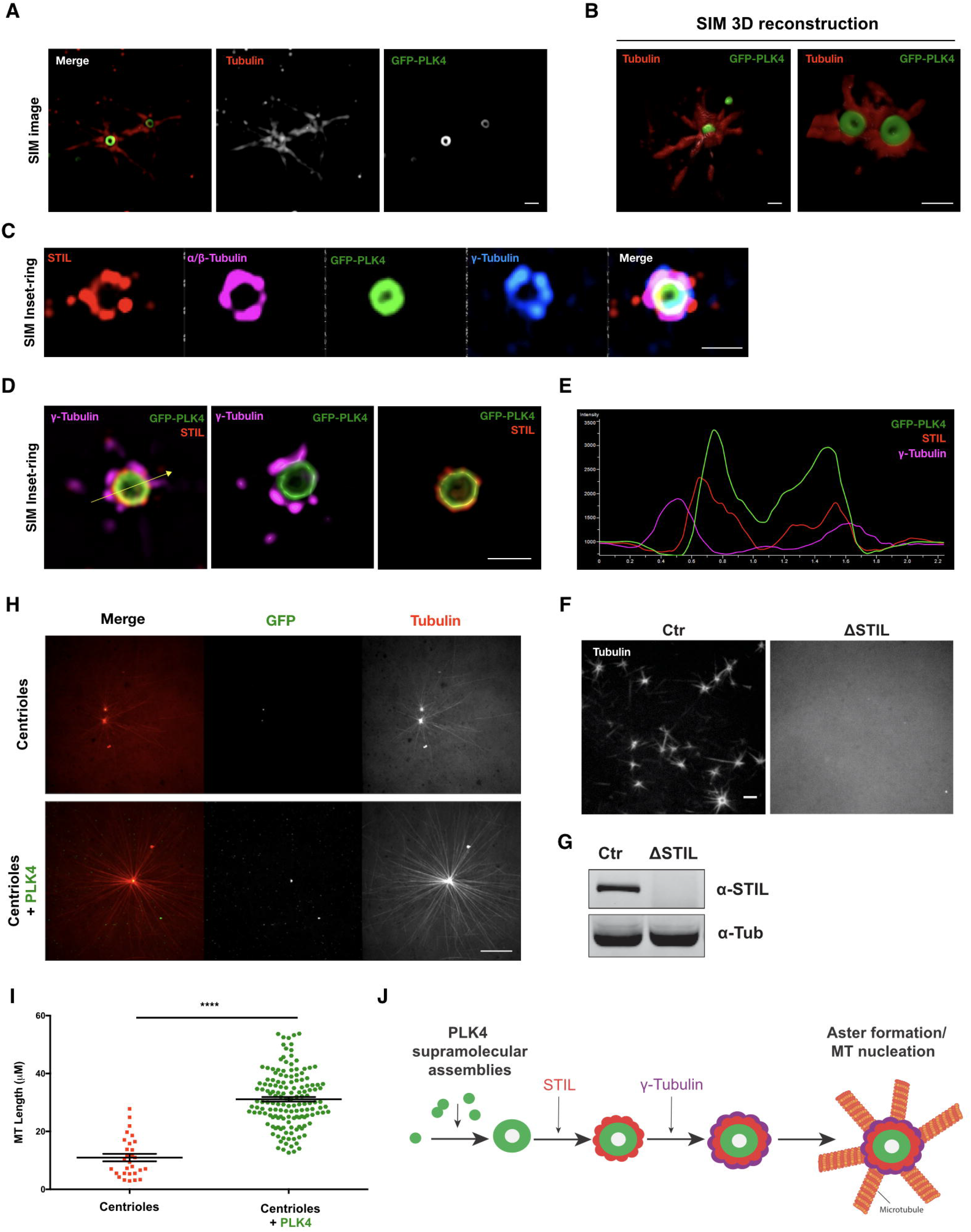
PLK4 supramolecular assemblies can recruit STIL and γ-tubulin in *Xenopus* released extracts, and are able to enhance centrosomal MT nucleation. (A) Representative images of 3D-SIM showing a ring-like structure of PLK4 MTOCs formed in calcium-released Xenopus extracts. α-tubulin and GFP-PLK4 are presented in red and green, respectively. Scale bars: 1 μm. (B) 3D-reconstitution of PLK4 asters (see Movie 2 and 3). (C) 3D-SIM images showing the co-localization of STIL (red), α-tubulin (magenta), GFP-PLK4 (green) and γ-tubulin (blue). Scale bars: 1 μm. (D) Representative SIM images showing the localization of GFP-PLK4 (green), STIL (red) and γ-tubulin (magenta) that co-localize with PLK4 supramolecular assemblies (see Movie 4). (E) Plots showing the intensity of each scored channels in (D). (F) Confocal images showing PLK4 induced-MTOCs in extracts (Control (Ctr)) and STIL depleted extract (ΔSTIL) using tubulin-rhodamine. (G) Western blot showing the depletion of STIL in the extracts used in (F). (H) PLK4 enhances MT nucleation. Confocal images showing MT nucleation using purified centrioles labelled with GFP-centrin incubated in *Xenopus* interphasic extract in the presence or absence of GFP-PLK4 (rhodamine-labelled tubulin (red); centriole and PLK4 (green)). Images were taken after 30 min incubation (see Movie 5 and 6). (I) Quantifications of MTs length (μM) visualized from the centrioles (GFP-centrin MTOCs) in presence or absence of GFP-PLK4. MTs were measured from 2 independent experiments, where 4 different MTOCs were analyzed. (N is the total number of MTs measured in presence of GFP-PLK4, N= 225). Scale bar: 5 μM. The statistical data are presented as ± s.d. ****p < 0.0001, (Mann-Whitney U).

We investigated the presence of STIL and γ-tubulin in PLK4 supramolecular assemblies. STIL is a well-known substrate of PLK4 and the formation of the complex PLK4-STIL is the first event that triggers centriole biogenesis (Loncarek and Bettencourt-Dias, 2017). γ-tubulin is a highly conserved protein, the major known MT nucleator, which is associated to all MTOCs studied so far (O’Toole et al., 2012; Teixido-Travesa et al., 2012). Most γ-tubulin in animal cells appears to exist as γ-TuRC that nucleates MTs (Wiese and Zheng, 1999). Interestingly, GCP6, one of the γ-TuRC members, is *a* PLK4 substrate (Bahtz et al., 2012; Martin et al., 2014).

We investigated the localization of STIL and γ-tubulin using immunofluorescence on fixed PLK4 MTOCs. We found that both STIL and γ-tubulin co-localize with PLK4 supramolecular assemblies in a spatially ordered manner (Fig. 4C and D). We observed PLK4 supramolecular assemblies in the centre, closely followed by a layer of STIL and then by γ-tubulin (Fig. 4D and Movie 4). The intensities of the three signals plotted together give a good insight of the close spatial relationship between the three molecules (Fig. 4E). We were unable to detect other centrosome proteins, due to the lack of specific antibodies against the *Xenopus* proteins. Importantly, depletion of STIL, prevents the formation of PLK4-induced MTOCs in extracts, suggesting that the same pathway is involved in triggering the formation of centrioles and acentriolar MTOCs (Fig. 4F and 4G) (Zitouni et al., 2016).

In summary, we have shown that *in vitro*, PLK4 self-assembles into supramolecular assemblies. When these supramolecular assemblies are added to extracts in interphase they can recruit STIL, γ-tubulin and α/β tubulin, forming a layered MTOC, similar to a centrosome. PLK4’s kinase activity is critical to form supramolecular assemblies and acentrosomal MTOCs (Fig. 1C, D, E and F), as well as centrioles (Moyer et al., 2015). Moreover, STIL is required for both PLK4-mediated centrosomal and acentrosomal MTOC formation, suggesting that both pathways use similar mechanisms. It is thus possible that even upon the presence of centrioles, PLK4 could promote the formation of supramolecular assemblies that concentrate components that are critical to form centrioles: including STIL, γ-tubulin and α/β tubulin. We have shown that both STIL and α/β tubulin bind PLK4. Since GCP6 (a known substrate of PLK4) is required for the recruitment of γ-TuRC to the centrosome, perhaps it can recruit γ-TuRC to PLK4-supramolecular assemblies (Bahtz et al., 2012; Oriolo et al., 2007; Teixido-Travesa et al., 2012). Additionally, CPAP, a binding partner of STIL known to be involved in MT stabilization, could indirectly enhance MT nucleation from PLK4 driven MTOCs (Sharma et al., 2016). PLK4 itself could also have a role in promoting further nucleation as it was recently proposed that MT stabilizers can promote MT nucleation in cells (Roostalu and Surrey, 2017). The authors proposed that MT stabilizers control the nucleation efficiency by stabilizing the MT centre or “nucleus”, either by providing a template for assembly or by promoting longitudinal or lateral tubulin-tubulin interactions.

Furthermore, it is also possible that the ability of PLK4 to bind MTs and to recruit STIL and γ-tubulin further promotes MT nucleation, even when centrosomes or other MTOCs are already present. To address this hypothesis, we used GFP-centrin purified centrioles from HeLa cells and incubated them with *Xenopus egg* extract. Shortly after addition to the interphasic *Xenopus egg* extract, the purified centrioles were able to recruit PCM components and nucleate MTs. However, when we added GFP-PLK4, we observed a very robust increase in MT nucleation capacity, MT elongation and a decrease in MT dynamics, suggesting their stabilization (Fig. 4H, 4I and Movie 5 and 6). We also observed the same effect in *Xenopus egg* extracts upon M-phase release (Fig. S3). These observations are very similar to the ones from Popov and colleagues with XMAP215, a processed MT polymerase that plays an important role in MT nucleation in addition to γ-tubulin (Popov et al., 2002). Altogether, our observations suggest a mechanism by which PLK4 promote MT nucleation in centrosomal and acentrosomal systems.

PLK4 forms supramolecular assemblies that recruit several important components in MT-nucleation, including gamma and α/β tubulin. We suggest this lowers the critical concentration of spontaneous MT nucleation leading to MTOC formation (Fig. 4J). Furthermore, the PLK4 supramolecular assemblies exhibit a layered organization, as it was shown to exist in the interphasic centrosome in animal cells (Lawo et al., 2012). The ability to mimic the layered centrosome in vitro opens up new ways of understanding PCM assembly. Future work aims at understanding how PLK4 supramolecular assemblies are formed, whether they form condensates, such as SPD5, and whether they are formed at the site of centriole birth, on mother centrioles.

## Materials and Methods

### PLK4 protein purification

Full-length *Xenopus* PLK4 gene lacking a stop codon was amplified by PCR and inserted into in-house-designed baculoviral expression plasmids (pOCC series) to generate the following construct: MBP-PreScission::PLK4::-mEGFP::PreScission-6xHis; The protein was expressed in SF+ insect cells and harvested 72 hr post infection. Cells were collected, washed, and resuspended in harvest buffer (50 mM Tris HCl, pH 7.4, 150 mM NaCl, 30 mM imidazole, 1% glycerol) + protease inhibitors (1 mM PMSF, 100 mM AEBSF, 0.08 mM Aprotinin, 5 mM Bestatin, 1.5 mM E-64, 2 mM Leupeptin, 1 mM Pepstatin A)(Calbiochem) and frozen in liquid nitrogen. The protein was purified using a two-step purification protocol described in Woodruff, J. B., & Hyman, A. A. (2015)*. PLK4 clarified lysate was incubated first with Ni-NTA agarose beads followed by a second incubation with amylose resin. The MBP and 6xHis tags were cleaved and PLK4 was eluted by overnight incubation with PreScission protease. PLK4 was concentrated with a 50K Amicon Ultra centrifugal concentrator units (Millipore), aliquoted and flash frozen in liquid nitrogen. The lysis and final buffer used contained 50 mM Tris-HCl, pH 7.4, 500 mM NaCl, 0.5 mM DTT, 1% glycerol, 0.1% CHAPS. The elution buffer from the Ni-NTA beads contained additional 250 mM imidazole (Woodruff and Hyman, 2015).

### *In vitro* PLK4 supramolecular assembly

PLK4 supramolecular assemblies were formed by adding purified GFP-PLK4 (1 μM) to the condensate buffer (150 mM NaCl, 25 mM Hepes (pH 7.4) and 1 mM DTT). PLK4 was incubated for 5 min and then imaged using a spinning disk CSU-X1 (Yokogawa) confocal scan head coupled to a Nikon Eclipse Ti-E and controlled using MetaMorph 7.5 (Molecular Devices). For tubulin recruitment, tubulin-labeled rhodamines TRITC (Cytoskeleton) (500 nM) were added to the condensate buffer. We used BRB80 added to the buffer as a control. The assay using PEG (9%) was performed as described in (Woodruff et al., 2017).

### Microtubule pelleting assay

Tubulin was polymerized into MTs stabilized with 20 μM taxol in BRB80 buffer (25 mM HEPES, pH 6.8, 2 mM MgCl_2_, 1 mM EGTA, 0.02% Tween 20 (v/v)), and quantified by absorbance measurements at 280 nm. Various concentrations of MTs were mixed with constant concentrations of PLK4 in BRB80 Buffer or vice-versa. Samples (final volume=40 μl) were allowed to equilibrate at 37°C for 55 minutes, centrifuged in Airfuge at 90 000 rpm for 30 minutes, and both the supernatant (S) and pellet (P) collected and resuspended in SDS sample buffer, and equal amounts of supernatant and pellet were run on 4–20% Tris-HCl gradient gels (Bio-Rad). Gels were stained with Coomassie Blue or used for western blot used for MT and PLK4 detection. Quantification of the relative amounts of PLK4 in supernatants and pellets was performed using ImageJ (National Institutes of Health, Bethesda, MD). The dissociation constants measured by MT co-sedimentation represent the average and propagated error from three separate experiments.

### *In vitro* PLK4 and microtubule bundling on confocal assay

Taxol-stabilized MTs seeds were incubated with GFP-PLK4 (1 μM) for 15 minutes at 37°C and then mounted in a slide and observed by confocal microscope. Taxol MT seeds were done as previously described (Honnappa et al., 2009). Briefly, lyophilized 1mg of tubulin (Cytoskeleton) is resuspended In 100 μl of BRB80 buffer, GTP and MgCl_2_ and incubated for 30 minutes at 37°C. After 20 minutes taxol is added to a final concentration of 20 μm. Stored at room temperature.

### Electron microscopy negative staining assay

For the electron microscopy assays, GFP-PLK4 or GFP-PLK4^AS^ were mixed with purified MTs. Incubation was performed at 37<u>º</u>C during 30 minutes. To study the effects induced by NAPP1, the inhibitor was added after the mixing. Samples were adhered to glow discharged copper 150 mesh grids coated with 1% (w/v) formvar (®Agar Scientific) in chloroform (®VWR) and carbon. Following attachment, samples were rinsed with distilled water and stained with 2% (w/v) uranyl acetate. Electron microscopy images were acquired on a Hitachi H-7650 operating at 100 keV equipped with a XR41M mid mount AMT digital camera.

### Preparation of *Xenopus* Egg Extracts and MTOC Formation Assay

MII-arrested and interphase egg extracts were prepared as previously described (Lorca et al., 2010; Zitouni et al., 2016). Purified PLK4 (0.675 nM) was added to 20 μl of CSF extracts and released into interphase using calcium (20 mM). MTOCs were analyzed by using rhodamine-labeled porcine tubulin (Cytoskeleton). The assays using GFP-centrin labeled centrosomes, centrosomes were added to interphasic extract or to the MII-arrested extract containing rhodamine-labeled tubulin in the presence or the absence of GFP-PLK4 (1 μM). These extracts are incubated for 30 min at 16ºC and visualized by confocal microscope. Prism (version 5.0c; GraphPad) was used for statistical analysis and plotting when was needed.

### Super resolution of PLK4 in *Xenopus* extracts assay

Structured Illumination Microscopy (SIM) of PLK4-GFP supramolecular assemblies was performed on N-SIM, Nikon Inc., equipped with Apo TIRF 100x NA 1.49 Plan Apo oil objective, back-illuminated EMCCD camera (Andor, DU897), and 405, 488, 561 and 640 nm excitation lasers. 100 nm Z sections were acquired in 3D SIM mode generating 15 images per plane, and reconstructed. XYZ corrections were applied using the signals of 100 nm multi-spectral fluorescent spheres (Invitrogen) included in the sample.

### Correlative light and electron microscopy

To correlate light and electron microscopy images of PLK4-GFP supramolecular assemblies, *in vitro* PLK4-GFP self-assembly reaction mixture was overlaid directly to the coverslips mounted in Attofluor Cell Chambers (Invitrogen; A7816) and kept at 37°. The coverslips contained previously sparsely seeded and fixed HeLa cells. HeLa cells served as landmarks for subsequent identification of target Plk4-GFP supramolecular assemblies during trimming, sectioning, and imaging on electron microscope. Plk4-GFP supramolecular assemblies were fixed in 2.5% glutaraldehyde, and immediately imaged on an inverted microscope (Eclipse Ti; Nikon, Tokyo, Japan) equipped with a spinning-disk confocal (CSUX Spinning Disk; Yokogawa Electric Corporation, Tokyo, Japan), back-illuminated 13 μm pixel EMCCD camera (Andor, DU888), 100x NA 1.42 Plan Apo objective lens with 1.5x magnifying tube lens, and a 2x lens in front of the confocal head. The position of Plk4-GFP supramolecular assemblies were recorded by acquiring a stack of 200 nm-thick Z sections in fluorescent mode and then in DIC (using Nikon DS-U3 camera). The position of the target Plk4-GFP supramolecular assemblies and fiducial cells on the coverslip was marked by a diamond scribe, as described earlier (Kong and Loncarek, 2015). After fixation, the samples were washed in PBS for 30 min, pre-stained with osmium tetroxide and uranyl acetate, dehydrated in ethanol, and then embedded in Embed 812 resin. 80 nm thick serial sections were sectioned, transferred onto the formvar coated copper slot grids, stained with uranyl acetate and lead citrate, and imaged using a transmission electron microscope (H-7650; Hitachi, Tokyo, Japan) operating at 80 kV. For the alignment of serial sections and for image analysis we used Photoshop (Adobe) and Fiji (NIH).

## Author Contributions

S.M.G purified the proteins, performed the microtubule assays, the EM and the SIM experiments with the participation of P.D, E.T., A.L, D.K and J.L. The CLEM experiments were performed by D.K and J.L. S.Z. performed the aster formation assays in extracts and the supramolecular assembly formation assays with the participation of B.R.G. and T.H. S.M.G and S.Z wrote the manuscript and made the figures. All authors read, discussed and approved the manuscript. S.M.G. and M.B.D. conceived the study and were in charge of overall direction and planning.

## Acknowledgements

We are thankful to Anna Akhmanova, Raquel Oliveira and Jeffrey B.Woodruff for reading and discussing the manuscript. S.M.G was funded by an EMBO Long term fellowship ALTF 1088-2009, a Marie curie Intra-European fellowship (#253373), a FCT post-doc fellowship. The collaboration with J.L. laboratory in the USA was financed by a Journal of Biologist travel grant. S.Z is funded by ERC grant ERC-COG-683258. Research in JL lab was supported by the Intramural Research Program of the NIH, National Cancer Institute, Center for Cancer Research. M.B-D. Laboratory is supported by an ERC grant ERC-COG-683258 and FCT-investigator to MBD.

